# Robust neural face identity codes in the Super-Recogniser brain

**DOI:** 10.64898/2026.06.22.733666

**Authors:** Martina Ventura, Tijl Grootswagers, Timothy Cottier, Manuel Varlet, James Dunn, David White, Genevieve L. Quek

**Affiliations:** The MARCS Institute for Brain, Behaviour and Development, Western Sydney University, Sydney, Australia; School of Computer, Data and Mathematical Sciences, Western Sydney University, Sydney, Australia; School of Psychology, Western Sydney University, Sydney, Australia; School of Psychology, UNSW Sydney, Kensington, Australia

## Abstract

Super-Recognisers show exceptional ability in face recognition, providing a natural model of how perceptual systems optimise for individuating visually similar stimuli in variable viewing conditions. However, the neural representations supporting this extreme perceptual expertise are unknown. Here, we tested whether Super-Recognisers (n = 23) differed from typical recognisers (n = 21) in the dimensional organisation of neural face identity coding. We recorded 64-channel electroencephalography while participants viewed random and rapidly-presented sequences containing 10 naturally varying images of 40 unfamiliar identities. Using time-resolved representational similarity analysis we measured the geometry of identity representations, their consistency across observers, and how clearly they specified face identity. Although neural expression of identity information was robust in both groups, we found three key differences between Super-Recognisers and typical recognisers. First, the geometry of face identity representations differed between groups. Second, Super-Recognisers showed greater inter-individual consistency in representational geometry. Third, Super-Recognisers’ neural signals discriminated between face identities more strongly than those of typical recognisers. Differences in the coding of broader face categories (sex, age, ethnicity) were notably weaker, suggesting that the observed group differences reflected fine-scale differences in identity coding rather than global reshaping of representational geometry. Strikingly, all three differences emerged within a common mid-latency interval (∼300-500ms), implicating higher-stages of face processing associated with representations that are sensitive to face familiarity and link between perceptual and semantic domains. Together, these findings indicate that individual differences in face recognition ability reflect higher-level differences in neural identity coding, rather than enhanced early sensory processing.

**Significance statement:** Super-Recognisers have an exceptional ability to recognise unfamiliar faces, yet the neural basis of this advantage has remained unclear. We find brain activity expresses the identity of unfamiliar faces more strongly and consistently in Super-Recognisers than typical recognisers. These advantages were specific to individual identity and could not be explained by enhanced encoding of broader facial categories such as sex, age, or ethnicity. Critically, group differences emerged during a mid-latency stage of processing (∼300–500 ms) associated with higher-level representations that support stable identity coding across variable images. The findings suggest that exceptional face recognition arises from more robust neural representations of individual identity rather than enhanced early visual processing, providing new insight into the neural basis of perceptual expertise.

## 1. Introduction

Face recognition is our primary means for identifying people in daily life. This fundamental brain function underpins everyday social behaviour and has profound implications in forensic and security settings where accurate person identification is paramount. Yet face recognition ability varies widely across the population: At one end of the spectrum, people with developmental prosopagnosia have difficulty recognising even close friends and family, where those at the other end – so called ‘Super-Recognisers’ (Russell et al., 2009) – show exceptional performance on even the most challenging face identity processing tasks (White & Burton, 2022). Because face recognition requires fine discrimination of similar visual patterns that can vary markedly in appearance across different encounters (Jenkins et al., 2011), Super-Recognisers provide a natural model of perceptual expertise that can reveal how brain systems organise to solve challenging visual computations.

Cognitive and behavioural studies have established that Super-Recognisers outperform typical recognisers across a range of face matching and memory tasks (Noyes et al., 2017; Russell et al., 2009; Bobak, Dowsett & Bate, 2016b; Davis et al., 2016; White, 2025), with advantages that extend to applied settings such as forensic identification and law enforcement (Bobak, Hancock, et al., 2016; Ramon et al., 2019; Mayer & Ramon, 2023; Bate, Portch & Mestry, 2021). Importantly, these advantages persist under challenging conditions, including long retention intervals (Davis et al., 2020) and when only limited facial information is available (Dunn et al., 2022), indicating that exceptional face recognition is robust across diverse task demands (White, 2025).

Here we examine how face identity is encoded in Super-Recogniser brain activity. Understanding how neural encoding of face identity supports face recognition is a key goal for neuroscience and cognitive psychology. One influential framework proposes that faces are represented within a multidimensional “face space” in which identities occupy positions defined by their values along multiple dimensions (Valentine, 1991, 2016). Evidence from macaque face patches shows that identity can be reconstructed from population responses distributed across these dimensions (Chang & Tsao, 2017), while human EEG and fMRI studies reveal similar identity-specifying structure in patterns of neural activity where greater representational distance corresponds to improved identity discrimination (Carlin & Kriegeskorte, 2017; Nemrodov et al., 2018; Dobs et al., 2019; Ambrus et al., 2019; Chauhan et al., 2020).

Although the structure of neural face representations has been extensively characterised at the group level, far less is known about how these representations vary across individuals. Previous EEG studies have shown that Super-Recognisers can be distinguished from typical recognisers from as early as 70 ms after stimulus onset, with peak differences around the N170 stage of face processing (Faghel-Soubeyrand et al., 2024; Rossion, 2002). Similarly, enhanced P1 amplitudes have been reported in Super-Recognisers (Belanova et al., 2018). However, these effects appear to generalise across both face and non-face stimuli, while more selective markers of face-identity processing remain largely comparable between groups (Nador et al., 2025). Together, these findings suggest that exceptional face recognition may partly reflect a broader perceptual advantage that extends beyond faces (Nador et al., 2025; Stehr et al., 2026).

Critically, existing neuroimaging studies have not examined whether the structure of identity representations varies as a function of face recognition ability. From a face-space perspective, enhanced performance could arise from a more robust or coherent representational geometry that better separates individual identities. Importantly, the timing of these differences may provide insight into their origin: early effects would support enhanced perceptual encoding accounts (Nador et al., 2025; Schroeger et al., 2023), whereas later effects would suggest that exceptional recognition reflects downstream refinement of identity representations beyond initial visual processing.

Here we used time-resolved neural decoding of EEG activity (Robinson, Quek, & Carlson, 2023) to characterise face identity representations in Super-Recognisers (N = 23) and typical recognisers (N = 21). We targeted neural identity codes that were robust to naturalistic variation in different encounters with the same face, by recording EEG while participants viewed a steam of 400 images, including 10 naturalistic images of 40 unfamiliar identities. We show that Super-Recognisers share a more consistent and more separable neural identity code than typical observers, with these differences emerging during a common mid-latency processing stage associated with higher-level identity representations (Deen et al., 2025). Importantly, these effects could not be explained by differences in the encoding of broader categorical face properties such as sex, age, and ethnicity (Dobs et al., 2019; Quek et al., 2021).

## 2. Results

### 2.1. Super-Recognisers share distinct neural codes for face identity

Does the dimensional structure of neural face identity coding differ between Super-Recognisers and typical recognisers? We addressed this question by quantifying the similarity of neural identity coding across individuals, using correlations between participant’s representational dissimilarity matrices (EEG RDMs) at each timepoint. This analysis captures the extent to which different observers distinguish between identities in a similar relational pattern. This analysis revealed a significant correspondence between the empirical similarity structure and the group model during a mid-latency interval of approximately 330-355ms (Figure 1A), indicating that identity representational geometry clustered according to recognition ability during this stage of processing.

**Figure 1.**
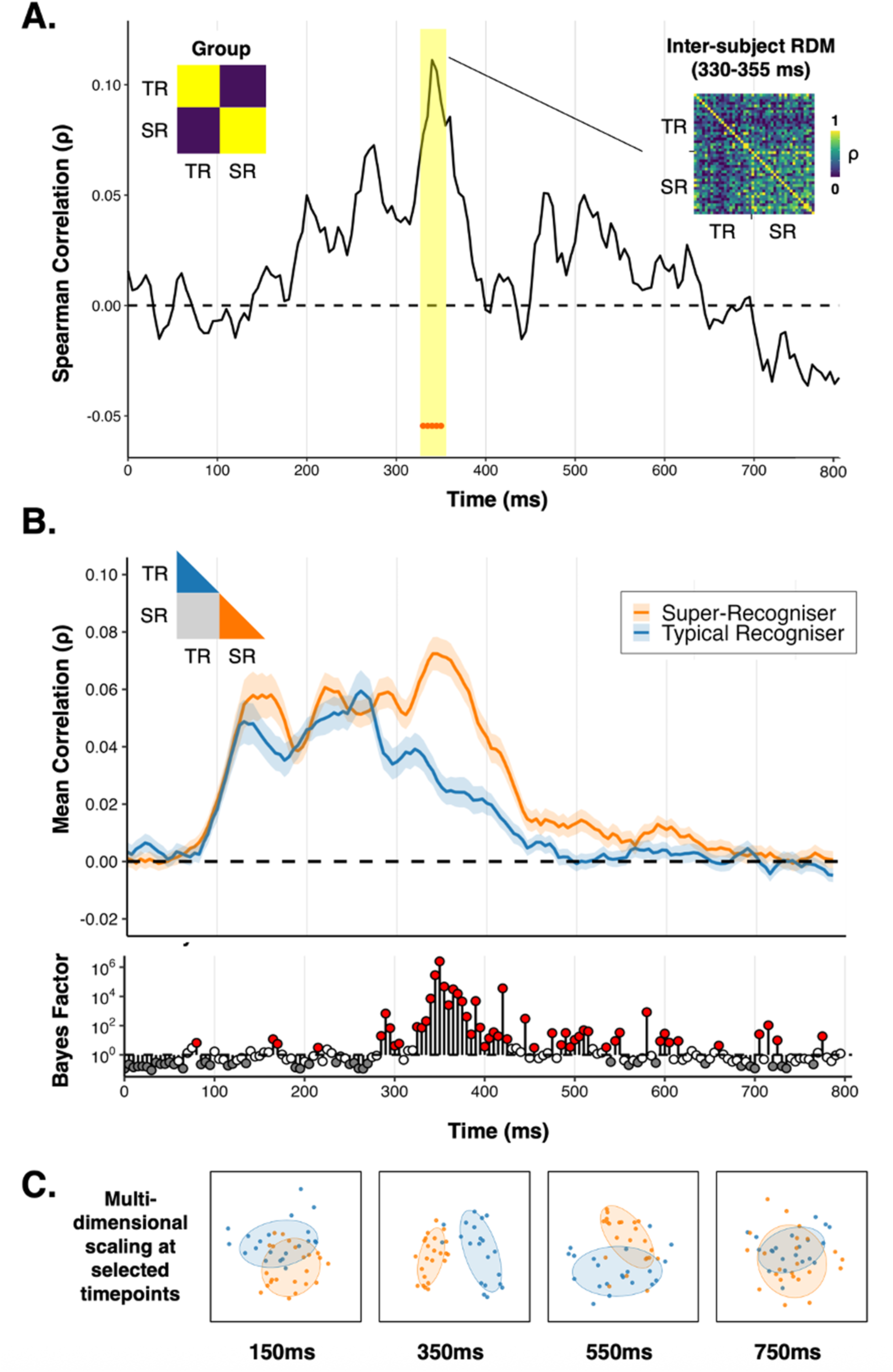
Super-Recognisers share distinct neural coding geometry. (A) Timecourse of between-group differences in the inter-subject similarity of neural face representations. The y-axis shows Spearman correlation (ρ) between the inter-subject similarity matrix and group-membership model that distinguishes participant pairs belonging to the same group from pairs belonging to different groups (inset left). Shaded region/red markers highlight the time window (330-355-ms) in which the correlation exceeded the permutation-based significance. The panel on the right shows the inter-subject representational similarity matrix averaged across the highlighted interval, illustrating stronger representational overlap in identity codes within the Super-Recogniser group. **(B)** Upper panel: Timecourse of inter-subject similarity in neural identity representations. Curves show the mean Spearman correlation between identity representational dissimilarity matrices among Super-Recognisers (SR: orange) and typical recognisers (TR: blue). Shaded regions represent ±1 SEM across participants. The dashed horizontal line indicates zero correlation. The inset illustrates the components of the inter-subject similarity matrix corresponding to SR–SR and TR–TR comparisons. Lower panel: bayesian evidence for differences in within-group similarity across time (SR *vs.* TR); red markers are time points with at least moderate evidence (i.e., BF>3). **(C)** Multidimensional scaling (MDS) visualisations of inter-subject representational similarity at representative time points. Points are individual participants; increased proximity indicates greater similarity in identity representational geometry. Ellipses capture the dispersion of participants in the Super-Recogniser (orange) and typical recogniser (blue) groups. Super-Recognisers cluster tightly together around ∼350ms, indicating greater convergence in these individuals’ identity representational structure.

We then generated an index of the similarity of neural coding geometry within each group by averaging the pairwise correlations between participants separately for Super-Recognisers and typical recognisers. As shown in Figure 1B, we found that identity representational geometries were reliably similar across individual participants within both groups (i.e., SR–SR and TR–TR), however, Super-Recognisers exhibited heightened between-participant consistency during a mid-latency interval of approximately 300-500ms. Although the magnitude of this group difference decreased beyond the 300-500ms interval, Super-Recognisers continued to exhibit higher inter-subject similarity into later stages of processing (approximately 450–700ms). Examples of the multidimensional scaling solutions for selected timepoints are shown in Figure 1C. Visual inspection of these solutions showed consistency with the group separation timecourse analysis, but representative timepoints showing the systematic changes in group separation are chosen for illustrative purposes. Together, these findings indicate that the geometry of identity representations in Super-Recognisers converges on a relatively consistent representational solution.

### 2.2. Enhanced separation of individual identities in Super-Recognisers’ neural signals

Our first analysis showed differences in the dimensional structure of neural face codes in Super-Recognisers compared to typical viewers in mid-to-late processing time window. Next, we compared the degree to which these representations discriminated individual face identities in Super-Recognisers and typical viewers. As shown in Figure 2A, despite substantial natural variability in both the low- and high-level properties of the face images (see Figure 4), both groups exhibited robust above-chance pairwise decoding of face identity over a sustained window between ∼100 and 500 ms. More importantly for our research questions, Bayesian comparison showed that Super-Recognisers exhibited higher identity decoding accuracies than typical recognisers during a mid-latency interval between approximately 310-360ms (Max BF = 8), with additional transient group differences re-emerging during later processing stages (550-770ms). Thus, while face identity information was reliably decodable in both groups, Super-Recognisers demonstrated enhanced identity separability during specific mid-to-late processing stages that corresponded with the epoch showing greatest convergence in Super-Recogniser’s face coding geometry.

**Figure 2.**
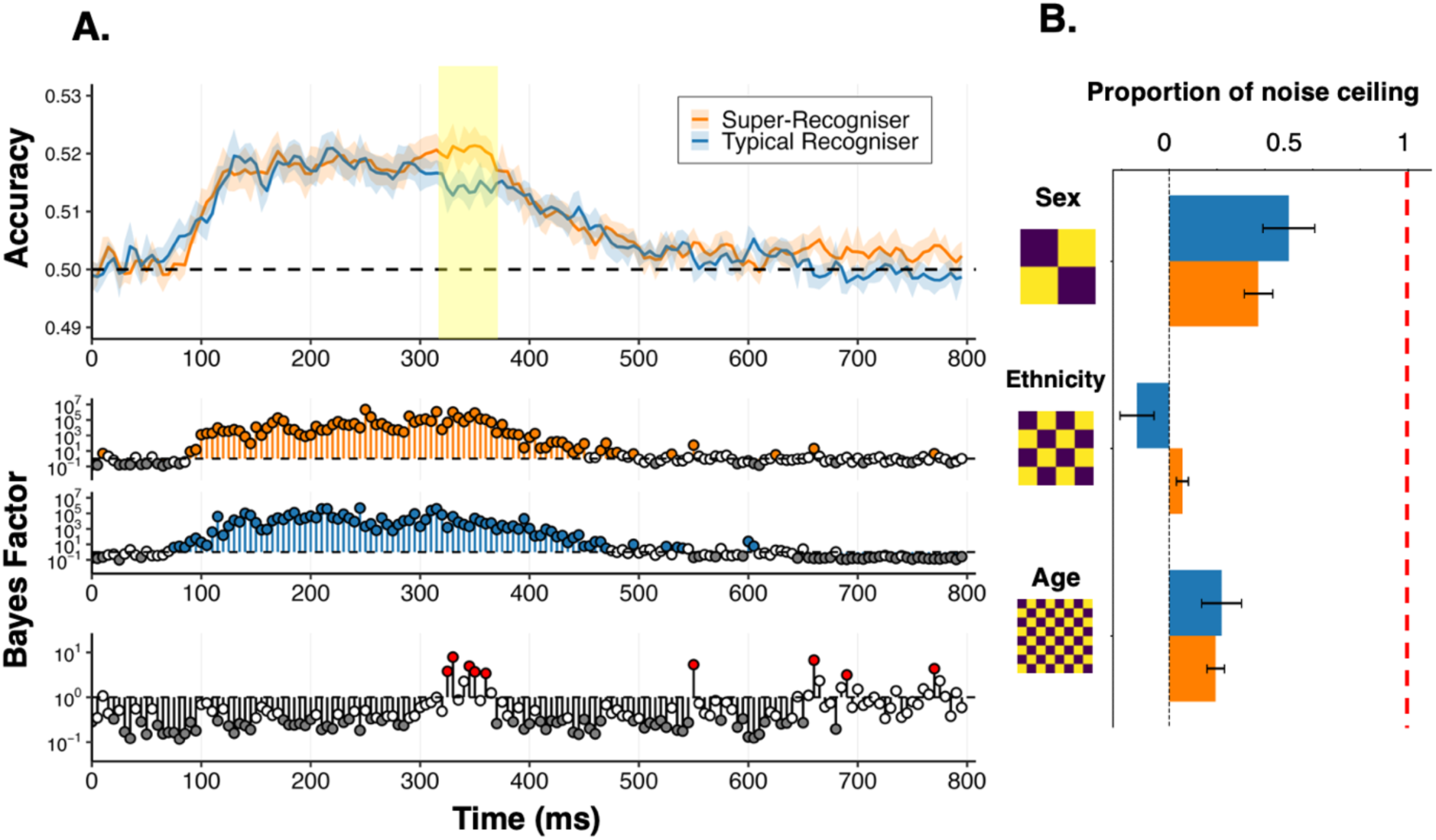
Identity decoding and inter-subject representational similarity. **(A)** Upper panel: Time-resolved pairwise decoding of the 40 unfamiliar face identities. Linear discriminant classifiers were trained to discriminate all identity pairs using leave-one-block-out cross-validation. Curves show decoding accuracy averaged across all identity pairs for Super-Recognisers (orange) and typical recognisers (blue); shaded regions are ±1 SEM. The dashed horizontal line denotes chance-level classifier performance (50%). Yellow shaded band highlights the time window in which Super-Recognisers exhibited higher decoding accuracy than typical recognisers (310-360ms). Middle panels: Bayesian evidence for above-chance pairwise identity decoding within each group. Lower panel: Bayesian evidence comparing decoding accuracy between Super-Recognisers and controls; red markers denoting time points with at least moderate evidence for a between-group difference (i.e., BF>3). **(B)** Correlations between categorical face aspect models and pairwise identity decoding averaged between 310-360ms, shown as a proportion of the noise ceiling estimate in each group. All model fits fell substantially below this ceiling, indicating that broad face categories are not sufficient to explain either the observed representational geometry of identity coding, nor the group-level difference, observed in this critical time window.

To verify that the stronger identity decoding observed for Super-Recognisers between 310-360 ms could not be accounted for by a group-level representational distinction in categorical face aspects (e.g., face-sex), we correlated participants’ neural RDMs averaged within this identified time window with theoretical models of face-age, face-sex, and face-ethnicity. Figure 2B shows the group mean model correlations expressed as a proportion of each group’s estimated noise ceiling – a value that reflects the maximum variance in the neural representational geometry that could be explained by *any* model given the consistency of the neural data across participants. For both groups, all model fits fell significantly below the noise ceiling (all *p* values < .001), indicating that none of the considered categorical dimensions fully accounted for the observed neural representational structure in the relevant time window. Even the strongest model correlation for both groups (face-sex) reached only around 50% of the noise ceiling, and was numerically higher for typical recognisers compared to Super-Recognisers, *t*(41.89) = 0.77, *p* = .444. Together, this shows that broad categorical organisation was not responsible for the enhanced identity decoding observed for Super-Recognisers during this time window, pointing instead to differences in fine-scale local organisation of face identity space.

### 2.3. Representational distinctions between Super-Recognisers and typical recognisers do not extend categorical face aspects

Although group-level distinctions in categorical face aspect coding do not account for the enhanced identity decoding for Super-Recognisers observed in Figure 2A, it could nevertheless be true that encoding of these face aspects differs generally between Super-Recognisers and controls. To assess this, our final analysis compared, over time, the degree to which neural signals in Super-Recognisers and typical viewers encoded the categorical face aspects. Figure 3 shows the time-resolved Spearman correlations between participants’ neural RDMs and theoretical models of face-sex, face-ethnicity, and face-age. Correlations with the face-sex model increased shortly after stimulus onset and remained positive across a broad interval (∼120-450ms) in both groups, with this correspondence notably stronger for typical viewers than Super-Recognisers during an earlier window between 200-300ms (Figure 3A). Information about face-age was also evident over a similar timecourse in the neural responses evoked in both groups (Figure 3C), however there was no indication that the strength of this representation differed for Super-Recognisers and controls. Finally, the face-ethnicity model was a comparatively weaker predictor of neural representational structure, with very modest positive correlations observed only for Super-Recognisers during a restricted mid-latency interval. Taken together, these results strongly imply that representational differences between Super-Recognisers and typical viewers in our study were not systematically related to differences in the broad category structure of face space.

**Figure 3.**
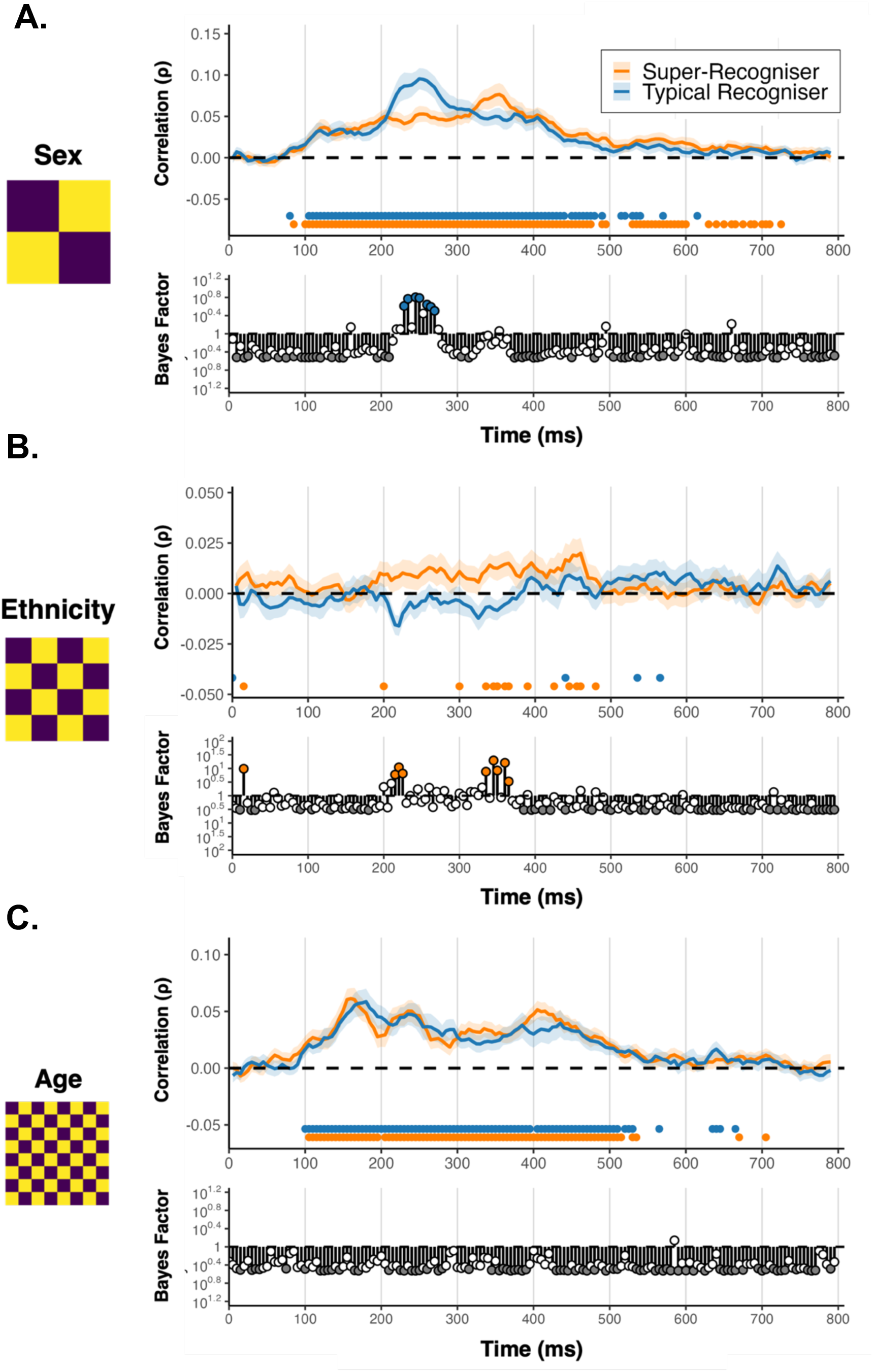
Time-resolved model correlations for sex (A), age (B) and ethnicity (C) information. For each representational model (left column), upper panels show Spearman correlations between pairwise identity decoding accuracy and the corresponding model predictions for Super-Recognisers (orange) and typical recognisers (blue). Shaded regions indicate ±1 SEM. Coloured markers below the curves denote time points with at least moderate evidence for above-zero model correlations within each group (i.e., BF > 3). Lower panels show Bayesian evidence comparing model correlations between groups, with coloured markers denoting time points with at least moderate evidence for a between-group difference (i.e., BF > 3)

## 3. Discussion

The present study examined how neural representations of unfamiliar face identities evolve over time and whether these representations differed between Super-Recognisers and typical recognisers. We found three key differences that distinguished brain activity patterns in Super-Recognisers. First, the representational geometry implied by brain activity differed between groups. Second, this coding geometry was more consistent across individual Super-Recognisers than across individuals with typical levels of face recognition ability. Third, the identity of a face was more reliably encoded in the brain activity of Super-Recognisers. Critically, all three of these group differences showed striking consistency in time course, emerging within a common mid-latency interval (∼300-500ms). These differences also appear to reflect coding of more fine-grained aspects of face identity that were not captured by broad category structures of the face space.

We studied identity representations under rapid serial presentation, conditions that minimised strategic or task-driven processing. Nevertheless, and despite images varying naturally in viewpoint, expression, lighting, and other image properties, identity information was robustly decodable in both Super-Recognisers and typical recognisers across extended intervals. Previous time-resolved neuroimaging studies have shown that identity information can be detected from neural responses to faces (Nemrodov et al., 2016, 2018; Ambrus et al., 2021), although these effects are typically stronger and more persistent for familiar faces (Barragan-Jason et al., 2015; Kovács et al., 2023). These findings indicate that stable neural identity codes can emerge under passive viewing conditions and across the type of image variability that characterises real-world face recognition (Jenkins et al., 2011; White et al., 2022).

Although identity information was reliably expressed in both groups, exceptional face recognition ability was associated with stronger and more consistent identity representations. Super-Recognisers exhibited greater identity separability together with increased inter-subject similarity of representational geometry. Thus, what differentiated the groups was not only the greater evidence of identity information carried by neural information, but also the way that information was organised.

This distinction is important for understanding how individual differences in face recognition arise. Face-space models conceptualise identities as positions within a multidimensional representational space whose geometry can be modified by experience (Valentine, 1991). Human fMRI and primate electrophysiology studies indicate that identity is represented as distributed patterns of distances between faces rather than discrete identity-specific units (Carlin & Kriegeskorte, 2017; Chang & Tsao, 2017). Behavioural studies further show that familiarity reshapes these relationships, increasing perceived similarity among images of the same individual while enhancing the separation between different identities (Chauhan et al., 2020; Kramer et al., 2025). Our findings extend this characterisation that face-space geometry is not fixed, but can be refined through experience and individual differences in face recognition ability. Further. because this relational identity structure was more consistent between individual Super-Recognisers than between typical recognisers, our data point to a common geometry that optimises face identity processing in the human brain.

Importantly, these group differences converged within a common mid-latency interval (∼300-500ms), with increased inter-subject similarity, enhanced identity discrimination, and stronger identity geometry, all emerging during this stage. The convergence of these independent measures suggests that individual differences in face recognition ability are expressed most strongly during later stages of identity processing rather than during initial perceptual encoding. Contemporary models propose that identity representations progressively dissociate from image-based similarity during this period, giving rise to more stable and abstract identity codes capable of linking to other person information (Wiese et al., 2024; Wang et al., 2022). Consistent with this view, activity within the 200-500ms interval has been linked to the strengthening of identity representations and increasingly sustained familiarity-related signals (Dalski et al., 2022).

The mid-latency interval showing most robust group differences fell between ERP components showing evidence of identity learning (N250) and links to semantic identity information (N400; see Wiese et al., 2024). N250 responses become more pronounced as initially unfamiliar faces become associated with stable identity representations (Tanaka et al., 2006; Kaufmann et al., 2009). More recently, Li and Li (2026) showed that the N250 emerges following visual familiarisation alone and is not further enhanced by identity labels or semantic information, suggesting that these processes reflect the establishment of stable visual memory traces rather than person knowledge per se. On the other hand, N400 responses show sensitivity to sematic relations between faces (e.g. Wiese & Schweinberger, 2015). How perceptual and semantic information are linked in neural face representations is not understood, but recent fMRI evidence points to a perirhinal cortex pathway linking to temporal pole regions as a key network in the formation of person identity representations (Deen et al., 2024).

Although the anatomical substrate underlying the extreme perceptual expertise of super-recognisers can not be inferred from our study, the timescale of our results do point to the role of of stable identity codes capable of linking person identity information. Rather than superior or faster extraction of visual information, our results are consistent with the enhanced formation of stable identity codes capable of specifying identity under substantial image variability. Because all faces in the present study were unfamiliar and viewed under passive conditions, the enhancement observed in Super-Recognisers implies that the mid-latency interval may support the formation of stable visual identity representations independently of stored person knowledge.

We also found that categorical information about faces was represented in neural responses. This finding is consistent with evidence suggesting that multiple facial dimensions contribute to the organisation of face representations. Behavioural and computational studies indicate that broad categorical attributes such as sex and age provide stable cues for structuring facial information (Valentine, 1991; Attard-Johnson et al., 2025; Rhodes et al., 2015). In the present study, these dimensions were represented in both groups and emerged relatively early, suggesting that they provide a common framework for face organisation. However, these dimensions accounted for a modest amount of variance in the overall identity geometry and did not to explain the enhanced identity codes observed in Super-Recognisers. In fact, the strongest categorical model (face-sex) was numerically larger in typical recognisers, indicating that superior performance cannot be attributed to enhanced encoding of coarse demographic information. Accordingly, exceptional face recognition here does not appear to arise from stronger encoding of broad categorical face information, but rather from enhanced refinement of identity-specific representations within a shared representational framework.

In conclusion, our results are consistent with representational accounts of perceptual expertise in which facial identity is encoded as structured geometry distributed across neural populations. Super-Recognisers exhibited stronger identity separation and greater inter-individual convergence and this was apparent within a mid-latency interval (∼300-400 ms). These differences point to more stable neural identity representations under image variability as underpinning individual differences in face recognition ability. However, broad similarities in other processing stages and in the overall category organisation suggest that exceptional face recognition reflects quantitative differences in the efficiency and precision of identity coding rather than qualitatively distinct processing mechanisms.

## 4. Materials and Methods

### 4.1. Participants

We recruited a total of 44 participants, consisting of 23 Super-Recognisers (18 F, Mean Age = 40, SD = 8.56) and 21 typical recognisers (13 F, Mean Age = 34, SD = 5.31). Super-Recognisers were recruited via the UNSW Face Registry, a database containing contact details of participants that have previously completed the following three standardised tests of face perception: The Cambridge Face Memory Test–Long Form (CFMT+; Duchaine & Nakayama et al., 2006), the Glasgow Face Matching Test–Short Form (GFMT; Burton et al., 2010), and the UNSW Face Test (Dunn et al., 2020). Super-Recognisers were individuals that achieved a composite *z*-score of at least 1.7 standard deviations above the mean on (Dunn et al., 2022). typical recognisers were recruited from the general population and were screened in advance of in-person testing using the Prosopagnosia-Index 20 (Shah et al., 2015), to exclude participants with self-reported face recognition difficulties. To ensure comparability with the Super-Recogniser sample, and to make sure they did not meet threshold to be considered a Super-Recogniser, all recruited typical recognisers underwent the same battery of behavioural face recognition assessments as described above (i.e., CFMT+, GFMT, UNSW Face Test). All participants were monetarily compensated for their participation.

4.2. *Stimuli*

We curated 400 face images consisting of 10 naturalistic images representing each of 40 unfamiliar identities, balanced by gender (male/female), age (older/younger) – where older refers to identities >65 years –, ethnicity (Caucasian/Hispanic). Images were sourced from Google; we deliberately allowed the face images to reflect natural variability that is observed in natural environments e.g., varied expressions, poses, head-angles, camera settings and lighting.

### 4.3. Procedure

Our experiment used a rapid serial visual presentation (RSVP) design that has been shown to support reliable time-resolved multivariate decoding of visual representations, even when neural responses to successive stimuli partially overlap; this reflects the capacity of the visual system to maintain multiple object representations simultaneously (Grootswagers et al., 2019). Participants viewed sequentially presented images at the center of a 24-inch ViewPixx monitor on a gray screen at a rate of 5 Hz (200ms per image, 5 images per second, no ISI between images). Each of the 400 images appeared exactly once per block in a randomized order, such that each sequence lasted 80 seconds. Participants underwent a total of 22 blocks, resulting in 220 presentations of each identity. To maintain attention, participants performed an orthogonal task, detecting randomly interposed upside-down face images by pressing a key. We used the inverted-target task in order to direct attention to the faces while not favouring any specific facial dimension. Feedback indicating the number of targets detected provided after each block (Figure 2). The presentation of stimuli and task control was managed using custom Python code and PsychoPy (Peirce et al., 2019). Throughout the experiment, we recorded continuous EEG data using a 64-channel BioSemi Active-Two electrode system with a sampling rate of 2048 Hz (BioSemi, Amsterdam, The Netherlands). Voltage offsets were maintained below 20mV. Electrode placement followed the 10/20 international standard (Oostenveld and Praamstra, 2001). The full EEG recording session lasted approximately 30 minutes, including breaks between sequences.

### 4.4. EEG pre-processing

Offline preprocessing was conducted using the Python MNE toolbox version 1.7 (Gramfort et al., 2014). The data were re-referenced to a common average and subjected to band-pass filtering with a high-pass filter set at 0.1 Hz and a low-pass filter at 100 Hz to eliminate slow drifts and high-frequency artifacts, including muscle noise. Epochs were extracted from -200 to 800ms relative to each stimulus onset with a baseline correction applied using the -200ms-0ms interval. Finally, the epoched data were downsampled to 200 Hz. No further preprocessing steps were applied; all channel voltages at every time point in the extracted epochs were utilized for the subsequent analyses. Because stimuli were presented at 5 Hz (200ms per image), epochs extending beyond stimulus offset necessarily contain overlapping neural responses from subsequent stimuli. However, the presentation order of the 400 images was fully randomised within each block, ensuring that responses to subsequent images (N+1, N+2) were not systematically related to the target image (N). As a result, any condition-specific signal that can be decoded at later latencies (e.g., 300-400ms) must be driven by the target stimulus rather than by overlapping activity from subsequent images. This approach has been validated in rapid serial visual presentation paradigms demonstrating reliable time-resolved decoding despite temporal overlap (e.g., Grootswagers et al., 2019, 2024; Robinson et al., 2019)

### 4.5. Analysis

#### 4.5.1. Identity-Level Representational Dissimilarity Matrices (RDMs)

To quantify the degree to which participants’ neural responses contained information about face identity that generalised across multiple images of the same person, we performed time-resolved pairwise decoding for epochs corresponding to the 40 unfamiliar identities (each of which was represented by 10 distinct images that varied substantially in viewpoint, facial expression, background, and lighting). For each unique pair of identities, we trained a Linear Discriminant Analysis (LDA) classifier to separate the two identities using a leave-one-block-out cross-validation procedure, in which each testing block functioned as the held-out test set exactly once. Classification performance was quantified using balanced accuracy (chance = 50%), and was run independent for each participant, at each time point, for all 780 unique identity pairs. For each participant, we assembled the full set of decoding accuracies for all identity pairs into a 40 × 40 symmetrical representational dissimilarity matrix (RDM) at each time point. Each cell of this RDM reflected the pairwise discriminability of two identities in that participant’s neural responses, with higher decoding accuracy interpreted as greater discriminability, i.e., identities that were more easily distinguished were positioned further apart in representational space. Constructing identity RDMs at the participant level allowed us to preserve the fine-grained structure of identity relationships over time and to directly compare representational geometries across individuals (Figure 1). Averaging across these pairwise classifications yielded a time-resolved measure of overall identity separability for each participant (Figure 2A).

#### 4.5.1. Inter-Subject Similarity Analysis of Identity RDMs

To assess whether identity representations were structured similarly across individuals, we directly compared identity-level RDMs between individual participants. Because each participant’s 40×40 matrix captures how that individual neurally distinguishes between every pair of identities, an inter-subject comparison enables us to quantify similarity in two individuals’ representational structure. At each time point, we used Spearman correlation to correlate pairwise identity discriminability values from one participant with those of every other participant. This yielded a time-resolved inter-subject similarity matrix, in which each cell reflected the degree to which two individuals distinguished between identities in a similar pattern. High similarity values indicate that identity pairs that were strongly separated in one participant’s neural responses were also strongly separated in the others’, whereas lower values indicate greater divergence in representational structure. We first asked whether this inter-subject similarity structure was organised by group membership. To do so, we constructed a theoretical group model in which pairs of participants belonging to the same group were assigned a value of 1, and pairs from different groups were assigned a value of 0. At each time point, the empirical inter-subject similarity matrix was correlated with this model (Spearman correlation), providing a time-resolved measure of the extent to which identity representations cluster according to group membership. We then examined the source of this group-level organisation by focusing on within-group similarity. Specifically, we quantified the similarity among Super-Recognisers (SR–SR) and among typical recognisers (TR–TR) separately across time. This allowed us to assess whether one group exhibited greater internal consistency in identity representational structure than the other.

#### 4.5.2. Predicting Identity-level RDMs with categorical face aspects

Notably, our pairwise identity decoding framework also allowed us to characterise the internal structure of identity representations and examine the degree to which identity signals carried information about categorical face aspects. For this purpose, we constructed three theoretical model RDMs representing face-sex, face-age, and face-ethnicity. In these binary models, each cell reflected whether the two identities belonged to the same (0) or different (1) category. Correlating the model RDMs with the observed neural RDMs enabled us to test whether these specific categorical aspects of faces were reflected in the neural responses evoked by viewing the 40 different identities, on the assumption that two identities differing along a given categorical dimension (e.g., different sexes) should evoke more discriminably neural responses than two identities that share that same dimension (e.g., same sex). Spearman correlations were obtained across the full timecourse (Figure 3), providing a time-resolved measure of how strongly each categorical dimension was reflected in participants’ evolving neural representations of faces.

To further examine model contributions during the period of strongest identity separation difference between the groups, model correlations were also computed within the time window identified in the decoding analysis (300-360ms). Within this interval, neural representational matrices were averaged over time for each participant, and model correlations were computed on these window-averaged representations. This yielded a single estimate, per participant and per model, of how much each social dimension accounted for the neural representational structure in this critical time window. To determine how much of the neural structure could in principle be explained by any model, we estimated a noise ceiling (Nili et al., 2014). For each participant, the window-averaged neural RDM was correlated with the average neural RDM of all other participants in the same group using a leave-one-out procedure, in which the target participant was excluded from the group average. This provided a conservative estimate of the maximum variance that any model could explain, given the consistency of the representational geometry across participants.

To further evaluate model performance relative to the explainable variance in the neural data, model correlations were also expressed as a proportion of the noise ceiling. For each participant, the full model correlation was divided by that participant’s own noise ceiling estimate. This analysis was conducted for the full correlations only.

### 4.6. Statistical inference

Statistical inference combined Bayesian analyses for time-resolved decoding measures (Morey et al., 2016; Teichmann et al., 2022) with permutation-based and frequentist approaches. We used the BayesFactor R package (Morey & Rouder, 2018) for all Bayesian analyses. Bayesian t-tests were implemented using a default Cauchy prior on standardized effect sizes (r = 0.707) (Wetzels et al., 2011). One-sided tests were used when a clear directional hypothesis was specified a priori, such as the prediction that Super-Recognisers would show stronger identity information than typical recognisers. In contrast, two-sided tests were used when no directional hypothesis was imposed, including analyses examining whether groups differed in the timing or magnitude of representational effects without assuming the direction of the difference. Bayes factors (BFs) were used to quantify the relative evidence for the alternative hypothesis compared with the null hypothesis. A BF indicates how much more likely the observed data are under the alternative hypothesis than under the null hypothesis. For example, a BF of 10 indicates that the data are ten times more likely under the alternative hypothesis than under the null. Here, BF values greater than 3 were interpreted as moderate evidence for the alternative hypothesis, values between 1 and 3 as anecdotal evidence for the alternative, values between 1/3 and 1 as anecdotal evidence for the null, and values smaller than 1/3 as moderate evidence for the null hypothesis.

Permutation testing was used to determine whether the similarity of identity representations across participants was structured by group membership. At each time point, we first computed the Spearman correlation between the empirical inter-subject similarity matrix and a theoretical group model that distinguished participant pairs belonging to the same group (SR–SR and TR–TR) from pairs belonging to different groups (SR–TR). This produced an observed timecourse describing how strongly inter-subject representational similarity aligned with group membership over time. To evaluate whether these correlations could arise by chance, a permutation procedure was used to generate a null distribution. Participant group labels were randomly shuffled while preserving the original number of SR and TR participants (5000 permutations). For each permutation, a new group model was constructed based on these shuffled labels, and the correlation between the empirical inter-subject similarity matrix and the permuted group model was recomputed across the entire timecourse. Repeating this procedure across many permutations yielded a distribution of correlation timecourses expected under the null hypothesis that group membership is unrelated to the structure of inter-subject representational similarity. To account for multiple comparisons across time, statistical significance was assessed using cluster-based permutation correction. First, a cluster-forming threshold was determined from the distribution of correlation values obtained across permutations. Time points in the observed correlation timecourse that exceeded this threshold were identified and grouped into consecutive temporal clusters. For each cluster, a cluster mass statistic was computed by summing the absolute correlation values across the time points belonging to that cluster. The same clustering procedure was then applied to each permutation timecourse, and for every permutation the largest cluster mass was retained. This produced a null distribution of maximum cluster masses expected under random group assignment. The statistical significance of each observed cluster was evaluated by comparing its cluster mass to this null distribution. Cluster-corrected p-values were computed as the proportion of permutations in which the maximum cluster mass was greater than or equal to the observed cluster mass. This quantifies how often an effect at least as large as the observed one would arise under random group assignment.

RSA was based on Spearman rank correlations, which provide a non-parametric measure of the correspondence between neural and model dissimilarity matrices. For the identity-level analyses, model correlations were computed between the neural representational dissimilarity matrices and the theoretical model matrices indexing sex, ethnicity, and age. These correlations were first computed across the full timecourse, yielding time-resolved estimates of how strongly each facial dimension explained variance in the neural representational geometry. Within each participant group, Bayesian one-sample *t*-tests were used to evaluate whether model correlations exceeded zero, and Bayesian independent-samples *t*-tests were used to assess group differences between Super-Recognisers and typical recognisers at each time point. To complement these time-resolved analyses, model correlations were also examined within a predefined temporal window corresponding to the period of strongest identity separation between groups, as identified by the decoding analysis (300-360ms). For this analysis, neural representational matrices were averaged across this interval for each participant, and model correlations were computed on the resulting window-averaged matrices. Within-group effects were assessed using two-tailed one-sample *t*-tests testing whether model correlations differed from zero, and group differences were assessed using independent-samples Welch *t*-tests. To estimate the amount of variance in the neural representational geometry that could be reliably explained by any model, a lower noise ceiling was computed separately within each participant group. For each participant, the window-averaged neural representational dissimilarity matrix was correlated with the average neural matrix of all remaining participants in the same group using a leave-one-out procedure, in which the target participant was excluded from the group average. This yielded one lower-noise-ceiling estimate per participant and provided a conservative estimate of the explainable variance in the neural data. Group differences in the lower noise ceiling were assessed using permutation testing. First, the mean lower noise ceiling was computed separately for Super-Recognisers and typical recognisers, and the observed group difference was defined as the difference between these two means (SR − TR). To generate a null distribution, group labels were randomly permuted across participants while preserving the original group sizes (5000 permutations). For each permutation, participants were reassigned to permuted SR and TR groups, the lower noise ceiling was recomputed within each group using the same leave-one-out procedure, and the resulting difference between group means was stored. Repeating this procedure generated a null distribution of group differences expected under the null hypothesis that group membership is unrelated to the consistency of the representational geometry. The permutation p-value was defined as the proportion of permuted group differences whose absolute value was greater than or equal to the absolute observed difference, thereby quantifying how often a group difference at least as large as the observed one would arise under random group assignment. To evaluate whether the tested models accounted for most of the reliably structured variance in the neural representational space, model correlations were compared with the estimated noise ceiling within each group using paired-samples t-tests. These tests assessed whether model fits differed significantly from the estimated ceiling of explainable variance. To control for multiple comparisons across models, p-values were corrected within each family of tests using the false discovery rate (Benjamini–Hochberg) procedure.

## Acknowledgements

We would like to sincerely thank all of the participants who generously contributed their time to this research. Their willingness to take part made this work possible. We are also grateful to Grace Wissman for her invaluable assistance with participant recruitment.

## Data and Code Availability

The data and analysis code supporting the findings of this study will be made publicly available upon publication.

